# Benchmarking the translational potential of AI-based drug-resistance prediction from *Mycobacterium tuberculosis* whole-genome sequencing data

**DOI:** 10.64898/2026.07.03.736369

**Authors:** Chang Liu, Haoyue Zhu, Pengfei Zhou, Nguyen Trung Thanh, Nguyen Quoc Dat, Ines Atmosukarto, Io Hong Cheong, Zisis Kozlakidis, Wiku Adisasmito, Xiaoqi Zheng, Hui Wang, Yang Yang

**Author notes:** These authors contributed equally to this work (Chang Liu, Haoyue Zhu, Pengfei Zhou).

## Abstract

**Background:** Tuberculosis, especially drug-resistant tuberculosis (DR-TB) including multidrug-resistant (MDR) and extensively drug-resistant (XDR) strains, remains a leading cause of infectious death worldwide. The rapid accumulation of whole-genome sequencing (WGS) data had spurred numerous computational methods for predicting antimicrobial resistance in *Mycobacterium tuberculosis*. However, heterogeneous datasets, preprocessing pipelines, and evaluation protocols have made fair comparisons impossible and have hindered clinical translation. A critical yet missing resource is a large-scale, unified benchmark to systematically assess and compare existing methods.

**Methods:** We curated an integrated MTB WGS–phenotypic drug susceptibility testing (pDST) dataset from three sources: the CRyPTIC dataset (Comprehensive Resistance Prediction for Tuberculosis: an International Consortium), a published multi-study compilation, and newly curated literature-derived datasets. The final benchmark contains 54,364 paired WGS–pDST records with broad geographic, lineage, and drug coverage. After harmonizing phenotypes and generating standardized variant features, we evaluated seven models (including classical machine learning and deep learning architectures) across 18 drug-level and six clinical resistance category prediction tasks.

**Results:** XGBoost achieved the highest mean drug-level AUPRC (0.674) and F1-score (0.620) and ranked first in AUPRC for 11 of 18 drugs, whereas WDNN achieved the highest mean AUROC. Random forest yielded the highest mean specificity (0.956) and accuracy (0.933), whereas logistic regression achieved the highest mean recall (0.774), highlighting distinct clinical trade-offs. Drug-level difficulty was highly heterogeneous: rifampicin and isoniazid were predicted robustly, whereas bedaquiline, delamanid, linezolid, and clofazimine remained persistently difficult. In clinical resistance category evaluation, RR-TB, MDR-TB, and pan-susceptibility were well predicted, but XDR-TB and other resistance categories constituted major bottlenecks.

**Conclusions:** Under the largest unified benchmark to date, classical machine-learning methods, particularly XGBoost, provided the strongest precision–recall and F1 performance overall, while neural models remained competitive by AUROC. Emerging drugs (bedaquiline, delamanid, linezolid, clofazimine) and XDR cases remain persistently difficult to predict, identifying key bottlenecks for future method development. This benchmark can serve as a community standard for evaluating MTB resistance prediction and the provided evaluation pipeline offers an actionable baseline for regulatory qualification and clinical decision support system validation, accelerating the translation of WGS-based resistance prediction into practice.

## 1 Background

Tuberculosis (TB) remains a leading cause of infectious-disease mortality worldwide, and drug-resistant TB (DR-TB), including multidrug-resistant (MDR) and extensively drug-resistant (XDR) strains, continues to undermine global control efforts. Rapid and reliable drug susceptibility testing (DST) is critical for guiding effective treatment, as inappropriate regimens accelerate the emergence and spread of resistance. Phenotypic DST (pDST) is the clinical gold standard, but it’s time-consuming, requiring up to eight weeks, creating a pressing need for molecular and computational approaches that can deliver earlier, clinically actionable predictions without sacrificing reliability [1, 2].

Whole-genome sequencing (WGS) offers a mechanistically informative route for resistance prediction, capturing known resistance mutations and broader genomic context. The WHO mutation catalogue and large WGS–pDST resources have established that WGS data can predict resistance to several first-line and second-line anti-TB drugs with high accuracy, provided that variants, phenotypes, and interpretive rules are consistently defined [3–5]. In parallel, a growing number of machine-learning (ML) and deep-learning (DL) methods have been proposed to learn resistance patterns directly from genomic features, including models specifically designed for multidrug resistance prediction [6, 7]. These advances make AI-assisted resistance prediction plausible, but they also make fair comparison among methods difficult, directly hindering clinical translation.

The root cause of this difficulty is fragmentation. Existing studies differ widely in isolate collections, pDST protocols, drug panels, variant-calling pipelines, feature encodings, data splits, and performance metrics. Fragmented datasets can overstate robustness; inconsistent preprocessing changes the effective input representation; and accuracy alone can be misleading in highly imbalanced resistance tasks. Moreover, geographic and lineage imbalances cause models that perform well under random splits to degrade in real-world deployment. Recent attempts have begun to address these issues. TB-bench focus exclusively on second-line drugs [8], and Big-TB primarily evaluate large language models on moderate-scale data [9]; however, both leave critical gaps unresolved due to their restricted scope. What remains truly missing is a large-scale, peer-reviewed benchmark that seamlessly integrates first-line and second-line drugs, encompasses broad geographic and lineage diversity, and provides fully standardized evaluation protocols from drug-level classification to clinical resistance categories in a single, reproducible framework.

To fill these gaps, we present four connected contributions: (1) we curated the largest integrated MTB WGS–pDST benchmark resources to date, combining CRyPTIC dataset, large published compilations, and newly curated literature-derived datasets, totaling 54,364 paired records with broad geographic, lineage, and drug coverage. (2) we established a unified preprocessing workflow that harmonises phenotypes, generates standardised variant features , and implements reproducible data splits. (3) we systematically benchmarked seven AI/ML approaches spanning linear models, tree-based methods, multilayer perceptrons, wide-and-deep networks, convolutional architectures, and published deep-learning models. (4) we evaluated translational aspects that matter for clinical deployment, including performance across 18 drug-level prediction and six clinical resistance category prediction tasks under lineage-stratified settings. In this study, we propose MTB-DRBench, a large-scale, clinically oriented benchmark for WGS-based MTB drug-resistance prediction under standardised data and evaluation protocols. Rather than asking only which model achieves the highest average accuracy, the benchmark identifies where models are robust, where they consistently fail, and which sensitivity–specificity trade-offs must be resolved before WGS-based AI can serve as clinical decision support. Importantly, the benchmark reveals that classical ML methods (particularly XGBoost) outperform more complex deep-learning architectures across most settings, and that emerging drugs (bedaquiline, delamanid, linezolid, clofazimine) together with XDR cases remain persistently difficult, highlighting key bottlenecks for future method development.

## 2 Methods

### 2.1 Data sources

We integrated three sources to construct the benchmark dataset, totalling 54,364 paired WGS-pDST records with broad geographic, lineage, and drug coverage.

#### CRyPTIC

The dataset curated by Comprehensive Resistance Prediction for Tuberculosis: an International Consortium (CRyPTIC) includes *Mycobacterium tuberculosis* isolates with matched phenotypic DST and whole-genome sequencing data from multiple countries. We used the public release after applying the benchmark-specific deduplication, phenotype harmonization, and feature-generation steps described in [5].

#### Published multi-study compilation

This includes samples from PATRIC, ReSeqTB, Coll et al., Hicks et al., Dheda et al., Zignol et al., Wollenberg et al., and Phelan et al., providing additional phenotypic resistance profiles and geographic metadata [10–16].

#### Literature-derived data (2021–2024)

we searched PubMed, Web of Science, and Embase for articles containing keywords related to *M. tuberculosis*, phenotypic drug susceptibility testing, and whole-genome sequencing between January 1, 2021, and December 19, 2024 (Table 1). Of 854 initial records, 344 were screened after duplicate removal, and 24 studies provided paired WGS and pDST information. The PRISMA flow diagram is shown in Figure 1, and study characteristics are summarized in Table 2.

**Fig. 1.**
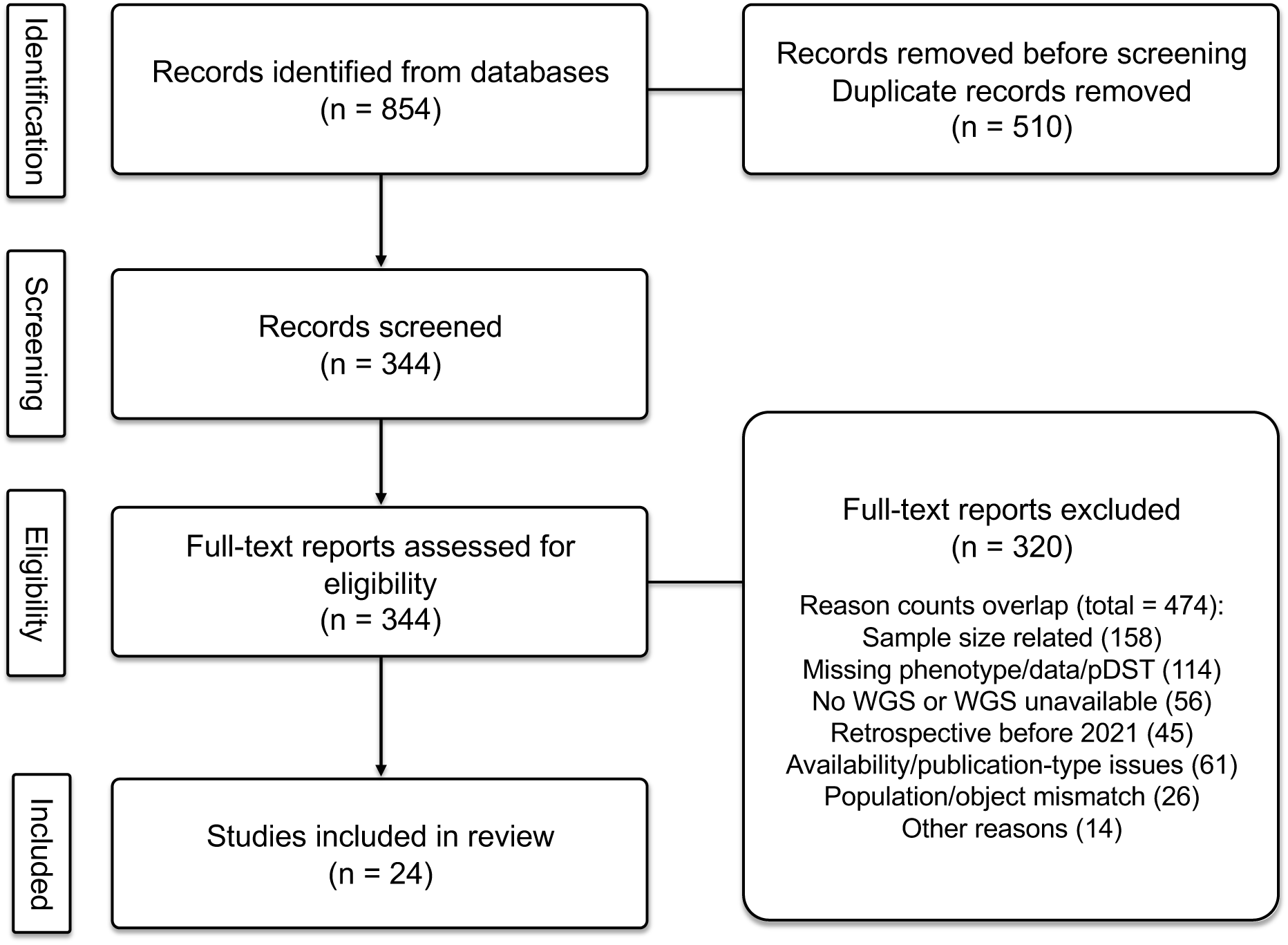
Screening process of the literature.

**Table 1.** Literature retrieval queries.

| Database | Query |
| --- | --- |
| PubMed | ("Tuberculosis"[Mesh] OR "MTB") AND ("Drug Resistance"[Mesh]) AND ("Microbial Sensitivity Tests"[Mesh] OR "DST" OR "drug sensitivity test") AND ("WGS" OR "whole genome sequencing") AND (2021:2024[pdat]) |
| Web of Science | ((TS=(MTB)) OR TS=(tuberculosis)) AND TS=(drug resistance) AND (TS=(DST) OR TS=(drug sensitivity test)) AND (TS=(WGS) OR TS=(whole genome sequencing)) |
| Embase | ('tuberculosis'/exp OR 'mtb:ab,ti') AND 'drug resistance'/exp AND ('microbial sensitivity test'/exp OR 'dst:ab,ti' OR 'drug sensitivity test'/exp) AND ('whole genome sequencing'/exp OR 'wgs:ab,ti') AND (2021:py OR 2022:py OR 2023:py OR 2024:py) |

**Table 2.**
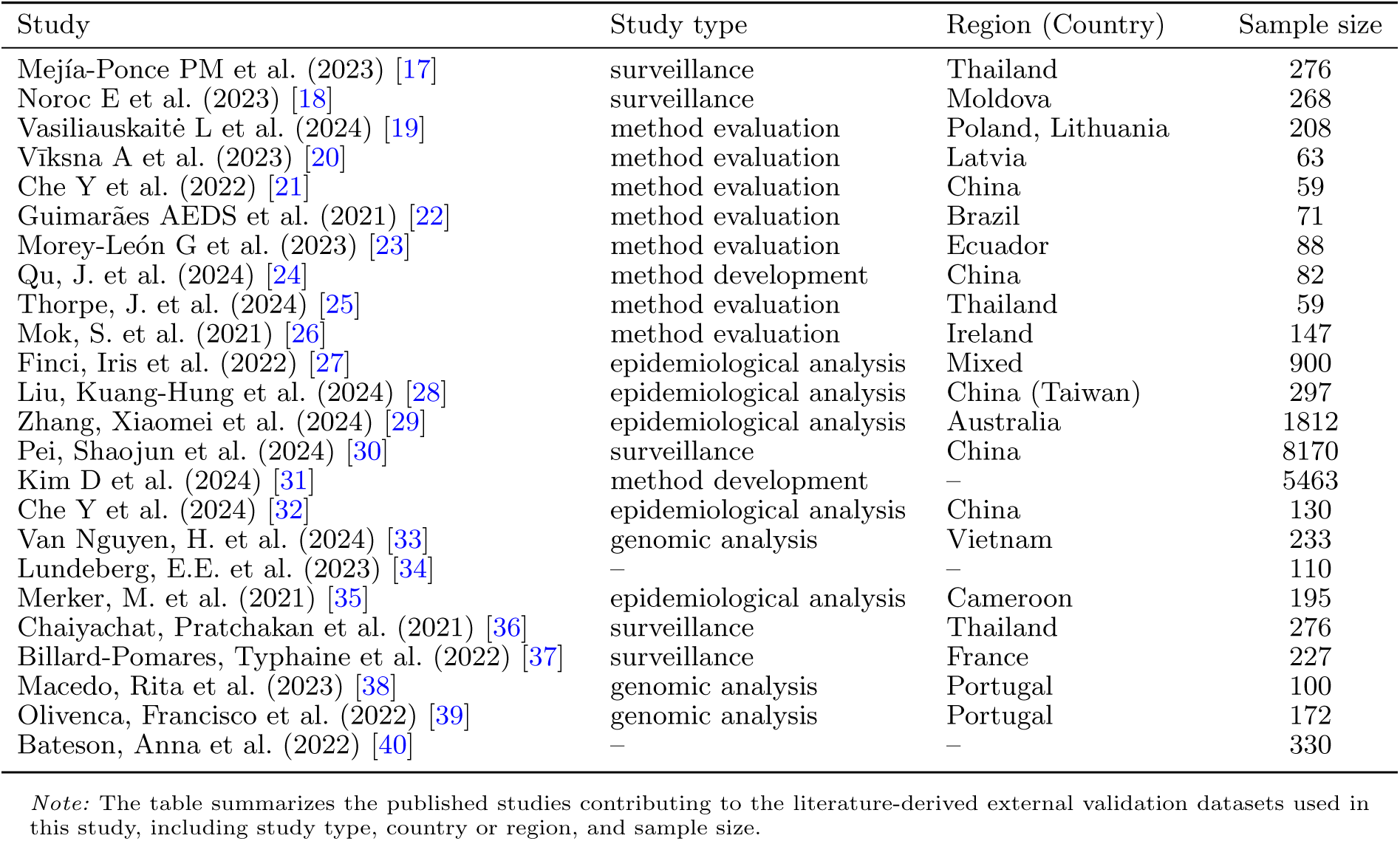
Literature-derived validation studies.

### 2.2 Phenotypic and genotypic quality

Phenotypes were harmonized to binary susceptible/resistant labels. Intermediate, indeterminate, contaminated, missing, or otherwise uninterpretable pDST results were excluded per drug rather than imputed. The names of drug were standardised, including harmonisation of common aliases such as capreomycin/CAP/CM, kanamycin/KAN/KM, and streptomycin/STM/SM. Duplicate isolates were resolved by retaining the most complete WGS–pDST profile; if discordant phenotype calls persisted, the isolate–drug pair was removed.

To audit pDST heterogeneity, each isolate–drug phenotype was assigned a method category: (1) standard culture-based pDST on Löwenstein–Jensen, Middlebrook 7H10, Middlebrook 7H11, or BACTEC MGIT (Becton Dickinson) using WHO DST guideline critical concentrations; (2) the same media or platforms but with outdated critical concentrations, or unreported concentration but cited WHO guidance; (3) non-standard methods or interpretive rules, with WHO critical-concentration testing and CRyPTIC-validated MIC interpretation treated as the standard references; (4) CRyPTIC-standard Thermo Fisher Scientific broth microdilution MIC with plate-specific epidemiological cutoffs; (5) insufficiently reported methods. Isolates were excluded if the WGS record lacked a valid sample identifier, could not be linked to pDST metadata, or failed bioinformatic quality control. Country, continent, and lineage annotations were harmonised for subgroup analyses.

### 2.3 Standardized preprocessing pipeline

All models were evaluated using an identical variant-derived feature representation to ensure that performance differences reflected model behavior rather than upstream differences. Public Illumina sequencing data were processed with Clockwork [41] for variant calling against the H37Rv reference genome (NCBI GenBank accession NC 000962.3). After sample-level and variant-level quality control, single-nucleotide variants and short insertions/deletions were annotated into standardised mutation features using Gnomonicus [42] and encoded as binary presence/absence variables. This mutation matrix was generated once before model fitting and reused across all benchmarked methods.

### 2.4 Benchmarked models

We evaluated seven representative AI/ML approaches, deliberately spanning classical machine learning and deep learning architectures:

#### Classical ML

logistic regression (LR), random forest (RF), extreme gradient boosting (XGB).

#### Deep learning

multilayer perceptron (MLP), wide-and-deep neural network (WDNN) [43], convolutional genome-wide prediction (CNNGWP) [44], and DeepAMR [7].

These models differ in capacity, inductive bias, and how they transform the shared variant matrix into predictions, as illustrated in Table 3. Each drug was treated as an independent binary classification task, and models were trained only on isolates with available phenotype for that drug.

**Table 3.** Overview of benchmarked ML models.

| Model | DL | Backbone / learner | Modeling assumptions | Parameterization / model size | Refs. |
| --- | --- | --- | --- | --- | --- |
| LR | × | Linear classifier | Linear additive effects | L2 penalty; $C = 0.9$ | [6, 45] |
| RF | × | Bagged trees | Non-linear interactions via bagging | 100 trees | [6, 46] |
| XGB | × | Boosted trees | Non-linear interactions via boosting | 1000 rounds; depth 6 | [47] |
| MLP | ✓ | Dense layers | Dense feature representation learning | 32–16 hidden units | [43, 48] |
| WDNN | ✓ | Wide & deep MLP | Wide-deep representation fusion | 64–64–64 deep branch | [43, 49] |
| CNNGWP | ✓ | 1D CNN + pooling | Local genomic-window patterns | 16 filters; kernel 10 | [44] |
| DeepAMR | ✓ | Encoder-decoder + single-task classifier | Latent mutation-profile representation | 128–64–16 encoder | [7] |

*Note:* DL, deep learning; $p$ , number of input variant features; $h_i$ , width of hidden layer $i$ . Trainable-parameter counts depend on the final feature matrix and selected hyperparameters.

### 2.5 Model training, validation, and reproducibility

Data were partitioned at the isolate level using the unique isolate identifier (uniqueid) after duplicate resolution. For each drug, we used nested five-fold cross-validation with random fold assignment stratified by the resistance label. In each outer fold, 80% of eligible isolates were assigned to training and 20% to testing. The outer test fold remained held out during model fitting, early stopping, and threshold specification. Within each outer training set, a further random five-fold split was used for early stopping where applicable. For neural models, early stopping monitored the inner-validation loss and was triggered when the loss did not decrease for 15 consecutive epochs. No spatial holdout split was performed. Lineage-specific robustness was assessed by calculating performance separately within the lineage groups represented in the outer test folds; lineages were not used as model input features.

Class imbalance was handled without resampling. LR and RF used balanced inverse-frequency class weights. For XGBoost, the positive-class weight was set separately in each fold to the number of susceptible training isolates divided by the number of resistant training isolates. Neural models were optimized with focal loss (*γ* = 2.0); its positive-class weight was updated for each batch as *α* = 1 − positive proportion and constrained to the interval [0.05, 0.95]. A fixed decision threshold of 0.5 was specified before outer-test evaluation and was not optimized by drug or model.

Metrics were calculated independently in each outer fold and averaged across the five folds for each drug. Overall model summaries were unweighted macro means across the 18 drug-specific tasks, and error bars represent the standard deviation across the five outer folds. Precision, recall, and F1-score were assigned zero when no positive predictions were made. AUROC was not imputed when an outer test fold contained only one class; a drug was omitted from an AUROC summary only if the metric was undefined in all five folds. All analyses used a master random seed of 2026 for Python, NumPy, scikit-learn, and PyTorch, with deterministic GPU operations enabled. Full software versions, model hyperparameters, and early-stopping settings are reported in Additional file 1.

### 2.6 Evaluation metrics and clinical resistance categories

Because resistance labels were strongly imbalanced for several drugs, we used the area under the pre-cision–recall curve (AUPRC) as the primary drug-level ranking metric. Secondary metrics included AUROC, F1-score, recall (sensitivity), specificity, precision, positive predictive value (PPV), negative predictive value (NPV), and accuracy. In addition to drug-level tasks, we further evaluated model performance on clinical resistance categories, defined according to WHO criteria:

#### Pan-susceptibility

susceptibility to all first-line anti-tuberculosis drugs (i.e. isoniazid (INH), rifampicin (RIF), ethambutol (EMB), and pyrazinamide (PZA)) [4].

#### HR-TB

INH resistance with RIF susceptibility.

#### RR-TB

RIF resistance with or without resistance to other drugs.

#### MDR-TB

resistance to at least both RIF and INH.

#### pre-XDR-TB

MDR/RR-TB with additional resistance to any fluoroquinolone (FQ).

#### XDR-TB

MDR/RR-TB with additional resistance to any FQ and at least one Group A drug, i.e. bedaquiline (BDQ) or linezolid (LZD) [2].

For each category, eligible samples were restricted to those with sufficient phenotypic information to assign a definitive 0/1 category label. Category-level predictions were derived from the model-predicted single-drug resistance outcomes using the same rule-based definitions, and performance metrics were then calculated separately for each category.

### 2.7 Use of large language models

During manuscript preparation, the authors used OpenAI Codex (GPT-5; OpenAI, accessed July 2026) to assist with English-language editing and improve clarity. The authors reviewed and revised all model-assisted text and take full responsibility for the content of the manuscript. The large language model was not used to generate data, perform statistical analyses, or determine the scientific conclusions.

## 3 Results

### 3.1 Benchmarked composition and data quality

The integrated MTB-DRBench combines three sources: the CRyPTIC public release, a large published multi-study compilation, and newly curated literature-derived records (2021-2024). Of total 54,364 isolates with paired WGS-pDST, 51,547 unique isolates have available origin and lineage metadata: 27,241 (52.9%) from CRyPTIC, 8,529 (16.5%) from the published multi-study compilation, and 15,777 (30.6%) from literature-derived records.

To our knowledge, this is the largest AI-ready unified MTB resistance benchmark, offering standardised data access, harmonised pDST labels, a unified genomic preprocessing workflow, and predefined benchmark splits (Table 4).

**Table 4.**
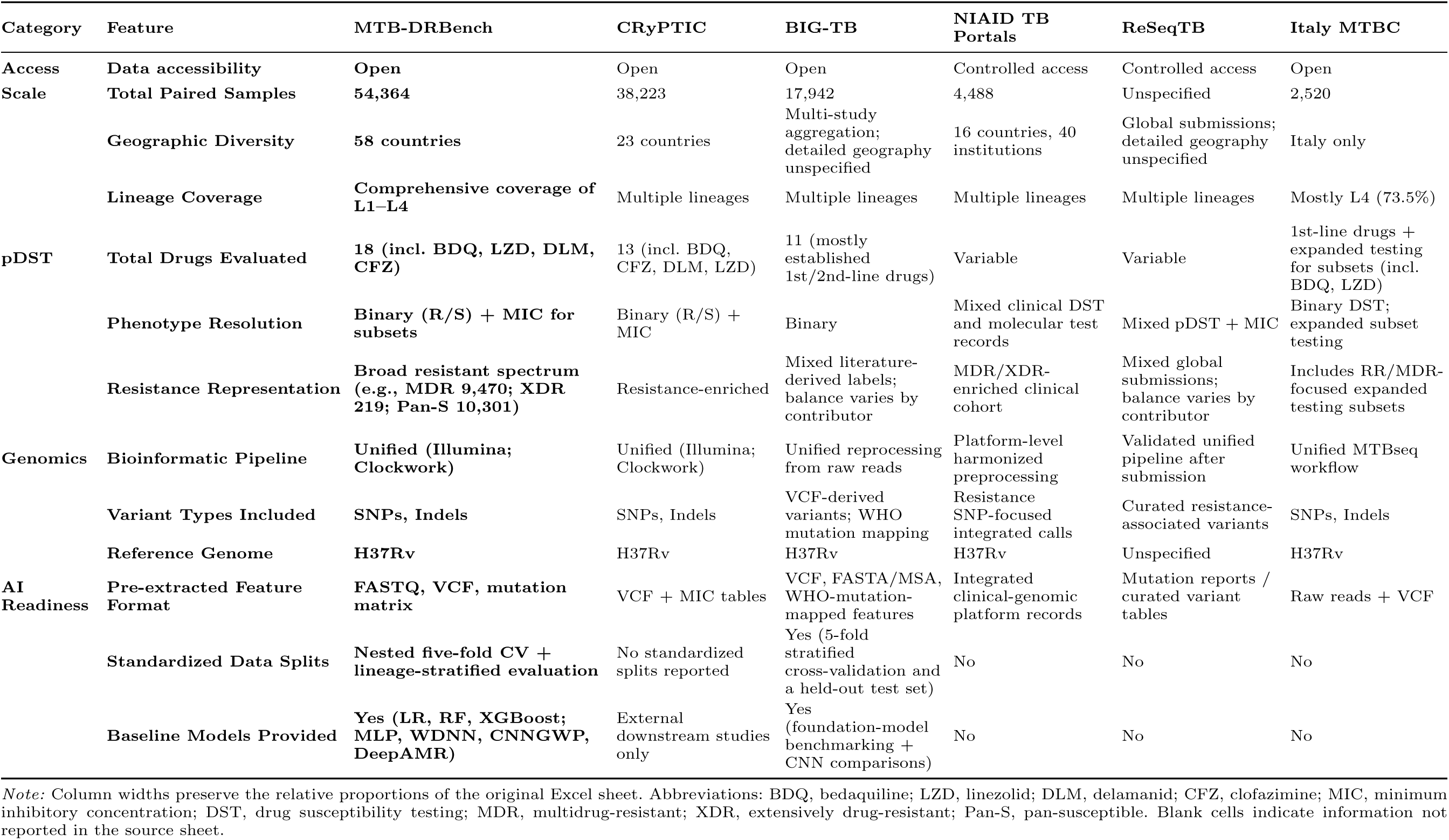
Comparison of the proposed MTB dataset with representative public TB resistance datasets and platforms.

Geographic and lineage representation is broad but uneven. Asia and Europe contribute the largest numbers of isolates, followed by Africa and South America, whereas North America and Oceania contribute smaller subsets. Lineage 4 and lineage 2 dominate, with lineage 3 and lineage 1 present at lower frequencies. This structure covers major global lineages but also motivates lineage-aware evaluation, because random splits can mask performance degradation in under-represented lineages.

We audited pDST method heterogeneity by assigning each isolate–drug phenotype to one of five categories (Table 5). Across 332,652 drug-level pDST observations, CRyPTIC-standard category 4 accounted for 159,540 (48%) records; category 2 records (outdated or incompletely reported WHO-based concentrations) 64,939 (19.5%); WHO-standard category 1 records 60,207 (18.1%); non-standard category 3 34,261 (10.3%); and unreported category 5 13,705 (4.1%). Category composition varies strongly by drug: emerging and repurposed drugs, such as BDQ, DLM, CFZ, LZD, LFX, and MFX, were dominated by category 4 MIC-derived calls, whereas PZA, RIF, CS, EMB, and INH have larger contributions from culture-based testing (category 1 or 2). This distribution makes the underlying method heterogeneity explicit for downstream interpretation.

**Fig. 2.**
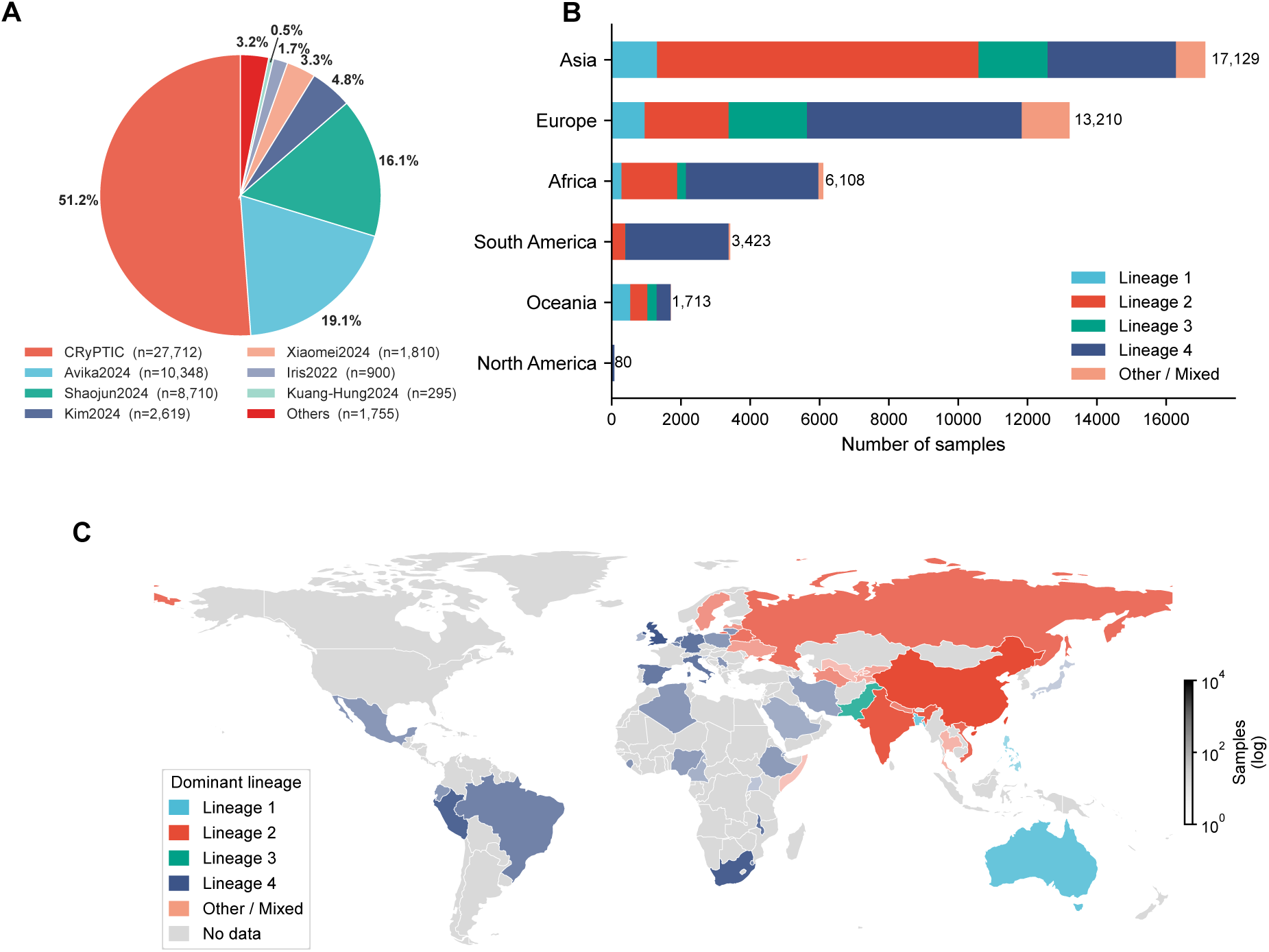
Composition and geographic distribution of the integrated MTB WGS–pDST benchmark. **(A)** Contribution of the top seven literature / projects sources to MTB-DRBench. Pie slices show the proportion of samples contributed by each source, with the remaining sources grouped as “Others”. **(B)** Geographic distribution of samples by continent and tuberculosis lineages. Stacked horizontal bars show the number of samples assigned to Lineage 1, Lineage 2, Lineage 3, Lineage 4, or other/mixed lineage groups. **(C)** Country-level global distribution of dominant tuberculosis lineage and sample availability. Countries are colored by their dominant lineage group, while gray indicates countries with no available samples; country-level sample counts are colored with a logarithmic scale.

**Table 5.**
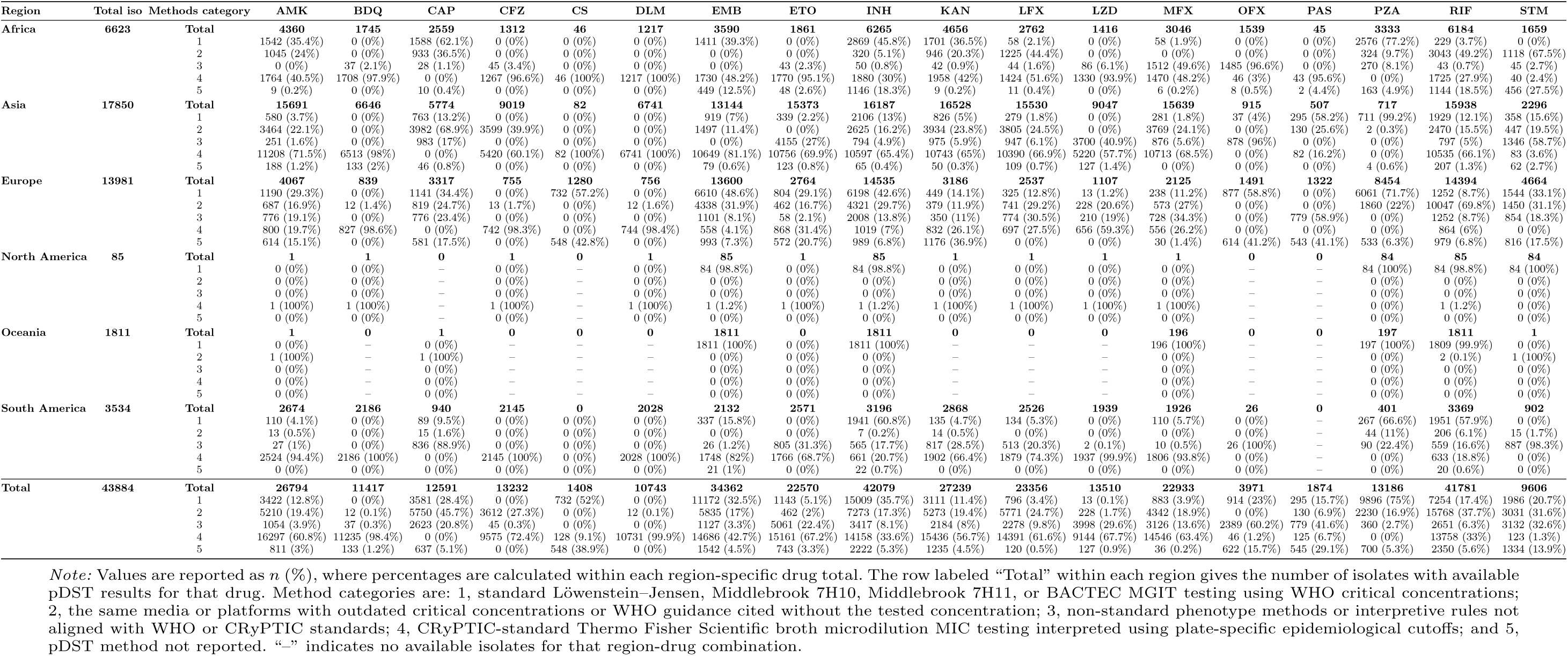
Regional distribution of pDST method categories across anti-tuberculosis drugs.

### 3.2 Drug-level prediction performance

All seven benchmarked models (Table 3) were trained and evaluated on the same variant feature matrix (binary presence/absence of Gnomonicus-annotated mutations) and the same data splits, ensuring that performance differences reflect model behavior rather than upstream preprocessing. For each of 18 drugs, we trained an independent binary classifier; the primary ranking metric was AUPRC due to severe class imbalance for several drugs. Specificity and accuracy remain clinically informative, but they were interpreted alongside recall and F1-score to avoid overvaluing models that mainly identify the susceptible majority class. This distinction was particularly important for BDQ, DLM, LZD, and CFZ, where resistant sample counts were low.

**Fig. 3.**
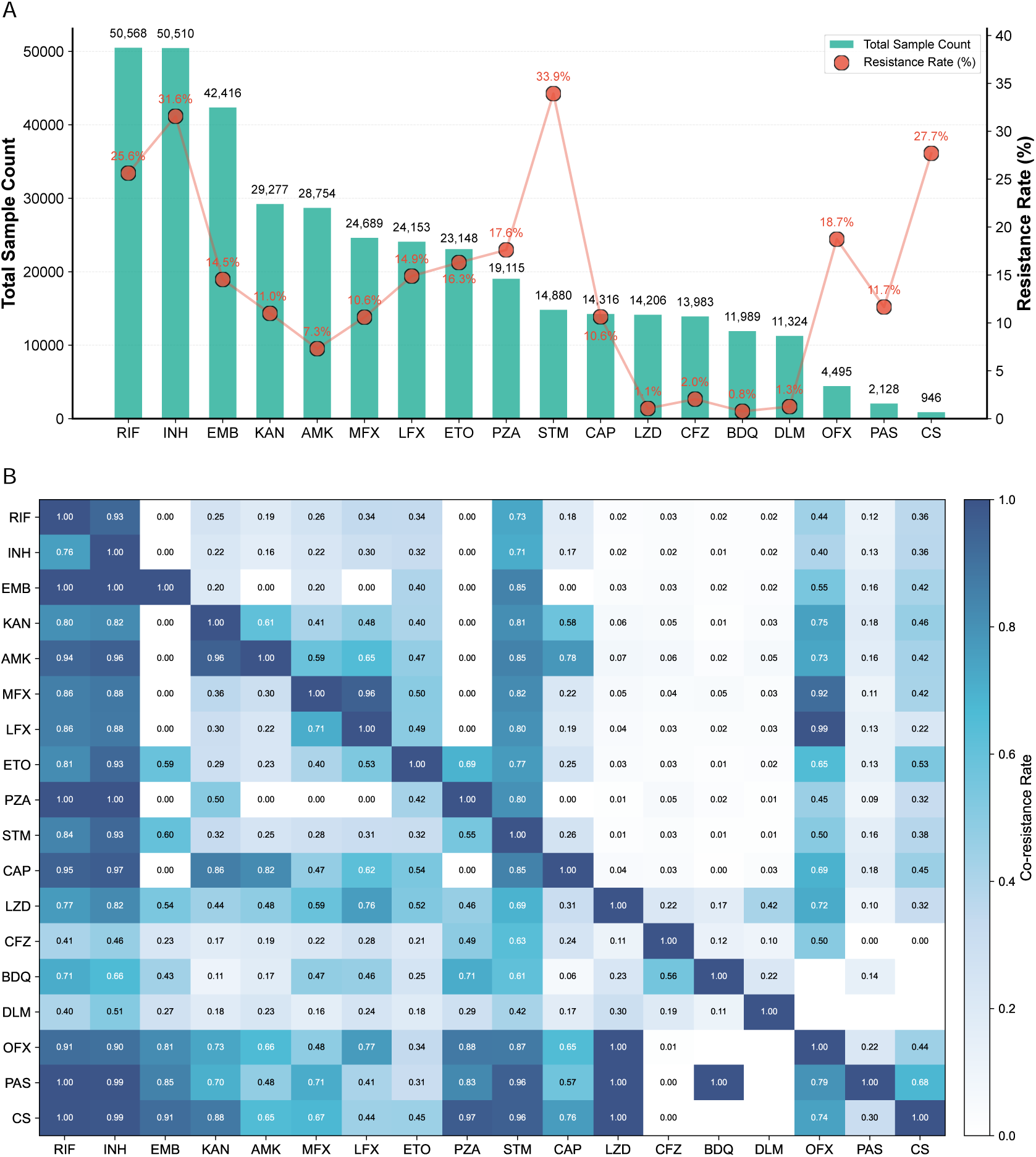
Drug-specific pDST coverage and resistance distribution in the benchmark. **(A)** Total sample counts and resistance rates for each anti-tuberculosis drug. Bars show the number of isolates with pDST information, while points and connecting lines show the corresponding resistance rate. Numeric labels above bars indicate total sample counts, and red labels indicate resistance percentages. **(B)** Pairwise co-resistance heatmap across drugs. Each cell shows the proportion of isolates resistant to the row drug that were also resistant to the column drug; darker blue indicates higher co-resistance. Diagonal values are 1.00 by definition, and blank cells indicate unavailable or undefined estimates.

#### Overall model ranking

XGBoost achieved the highest mean AUPRC across drugs (0.674 *±* 0.026) and the highest mean F1-score (0.620), and ranked first in AUPRC for 11 of 18 drugs (Figure 4). The next best models by mean AUPRC were MLP (0.657 *±* 0.029), WDNN (0.655 *±* 0.029), RF (0.646 *±* 0.022), and LR (0.644 *±* 0.030), indicating that the top-performing models were separated by modest differences relative to cross-validation variability. AUROC rankings showed a slightly different pattern, with WDNN (0.891 *±* 0.017) and MLP (0.887 *±* 0.017) achieving the highest mean AUROC, followed by LR (0.880 *±* 0.017), CNNGWP (0.879 *±* 0.018), RF (0.873 *±* 0.018), and XGBoost (0.871 *±* 0.018).

**Fig. 4.**
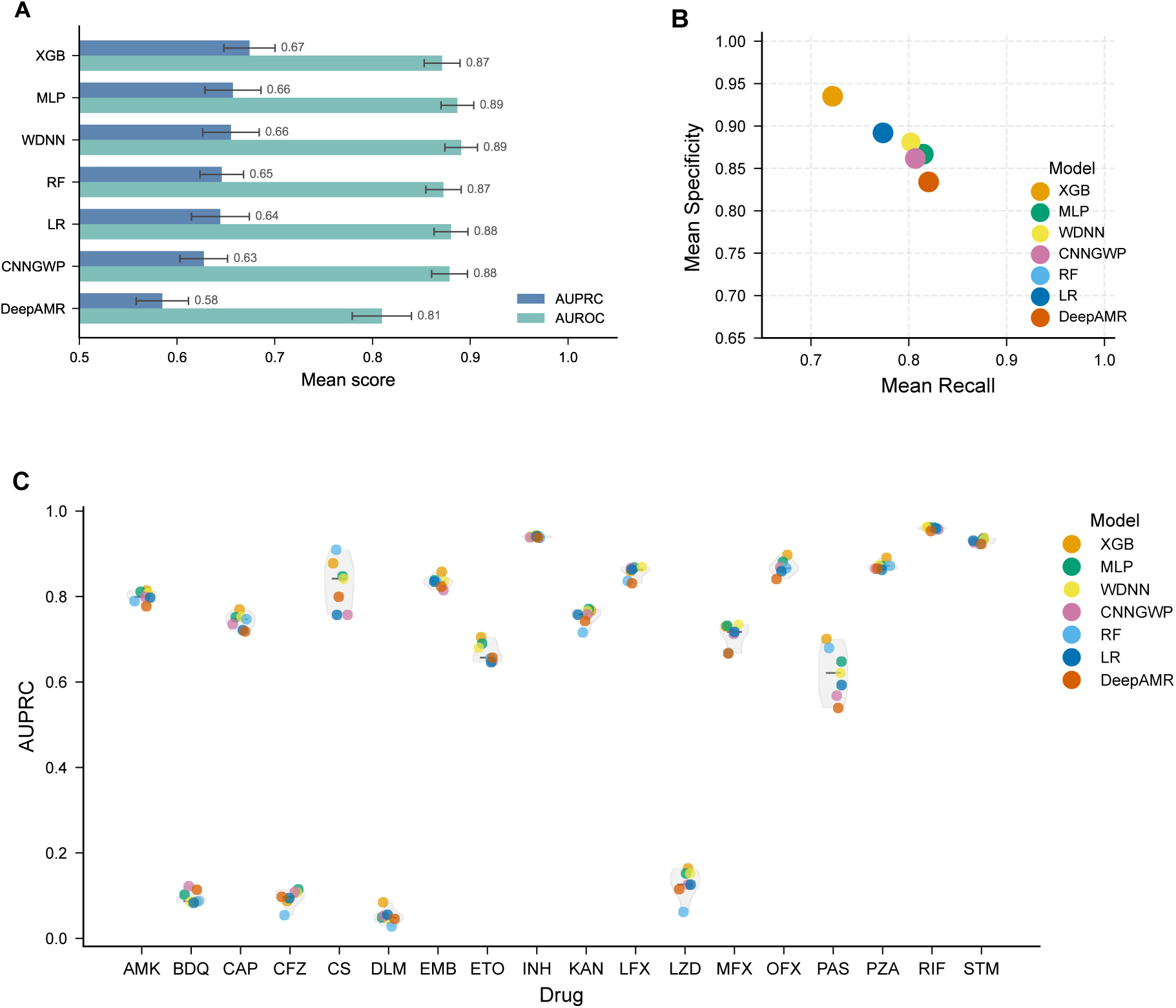
Overall drug-level model benchmark. **(A)** Mean AUROC and AUPRC of each machine learning model across drug-level binary prediction tasks. Bars indicate the mean performance across five cross-validation folds, and error bars indicate standard deviation. Models are ordered by mean AUPRC. **(B)** Mean recall and specificity of each model across drug-level tasks, showing the trade-off between sensitivity and false-positive control. **(C)** Distribution of drug-specific AUPRC values across models. Individual points represent model-level AUPRC values for each drug.

Thus, the optimal model depends on whether the clinical use case prioritises precision-recall performance under class imbalance, detecting resistance, avoiding false-resistant calls, or balancing both.

#### Drug difficulty is highly heterogeneous

Rifampicin and isoniazid were the easiest tasks (best AUPRC values of 0.963 *±* 0.005 and 0.943 *±* 0.002, respectively). In contrast, four drugs, i.e., CFZ (best AUPRC 0.108 *±* 0.026), DLM (0.085 *±* 0.051), BDQ (0.103 *±* 0.066), and LZD (0.192 *±* 0.090), were persistently difficult and showed larger fold-to-fold variability. These drugs had few resistant examples in the model-ready data (CFZ 275, DLM 142, BDQ 92, LZD 149), indicating that sample scarcity is a major driver. However, sample count alone does not fully explain difficulty: cycloserine and PAS also had limited resistant support but achieved higher best AUPRC (CS, 0.910 *±* 0.025; PAS, 0.701 *±* 0.072), suggesting that resistance mechanism complexity, label quality, and feature informativeness also contribute.

### 3.3 Lineage-aware and clinical category evaluation

#### Lineage-stratified robustness

When evaluated separately on each major lineage, all models showed incomplete robustness under distribution shift (Figure 5). XGBoost achieved the best F1-score in lineage 1, lineage 2, lineage 3, and lineage 4, whereas WDNN performed best in the mixed lineage group. For XGBoost, F1-score ranged from 0.436 in lineage 1 to 0.650 in lineage 2, demonstrating that high random-split performance does not guarantee generalisation across lineages. Fold-level AUROC/AUPRC summaries showed the same pattern of lineage-dependent variability. For XGBoost, AUPRC ranged from 0.514 *±* 0.122 in lineage 1 to 0.699 *±* 0.025 in lineage 2, with intermediate performance in lineage 3 (0.610 *±* 0.093) and lineage 4 (0.632 *±* 0.031). AUROC varied less in its mean values but still showed larger uncertainty in lineage 1 (0.810 *±* 0.096) than in lineage 2 (0.869 *±* 0.029), lineage 3 (0.858 *±* 0.059), and lineage 4 (0.847 *±* 0.042). This finding supports the need for lineage-aware validation before clinical deployment.

**Fig. 5.**
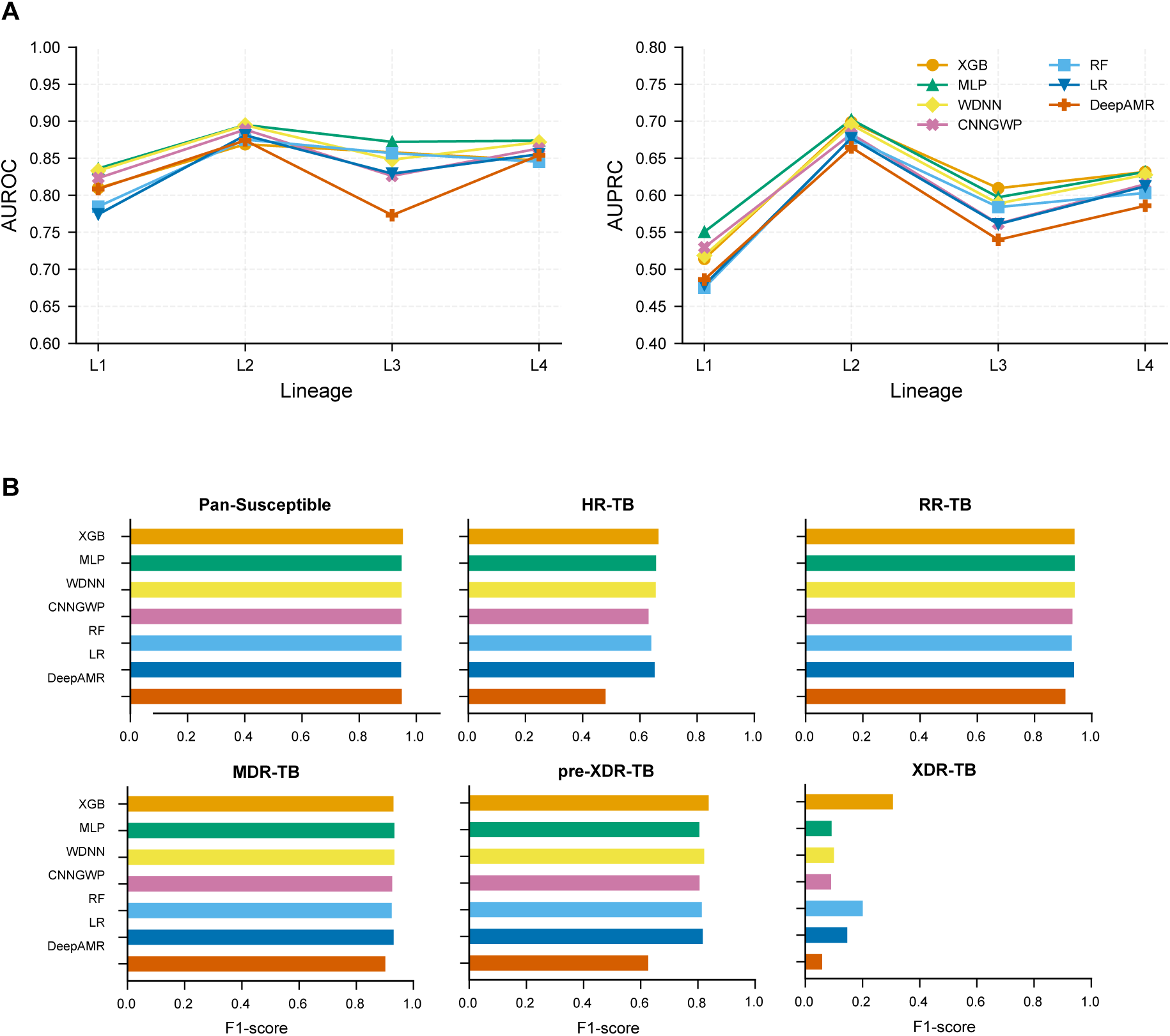
Drug difficulty and robustness across evaluation settings. **(A)** Lineage-stratified AUROC and AUPRC of each model across major tuberculosis lineage groups. Points indicate mean performance across cross-validation folds, and lines connect the same model across lineage groups to visualize robustness across lineage distributions. **(B)** Clinical resistance category performance by model. Horizontal bars show class-specific F1-scores for clinical resistance categories, including pan-susceptible, HR-TB, RR-TB, MDR-TB, pre-XDR-TB, and XDR-TB.

#### Clinical resistance categories

We evaluated six clinically defined categories using rule-based predictions derived from single-drug model outputs. Weighted F1-score was highest for RF (0.873), with XGBoost and MLP close behind (both approximately 0.870). Macro F1-score was highest for XGBoost (0.689). Common and high-signal categories were predicted well: best F1-score was 0.941 for RR-TB, 0.936 for MDR-TB, and 0.954 for pan-susceptible isolates. In contrast, **XDR-TB remained a major bottleneck** across all models, with best F1-scores of 0.330. This gap shows that clinical resistance category performance cannot be inferred from drug-level averages alone and that improving XDR-TB and emerging-drug resistance prediction should be a priority for future method development.

## 4 Discussion

This study establishes the largest unified benchmark to date for AI-based *Mycobacterium tuberculosis* drug resistance prediction from WGS data. By integrating 54,364 paired WGS-pDST records, standardising preprocessing and feature representation, and systematically evaluating seven models across 18 drugs and six clinical resistance categories, we provide a reproducible framework that directly addresses the fragmentation problem in the field.

### Model ranking depends on the clinical question

XGBoost achieved the highest mean AUPRC and F1-score, ranking first in AUPRC for 11 of 18 drugs, whereas WDNN achieved the highest mean AUROC. RF yielded the highest specificity and accuracy, whereas LR achieved the highest recall. These are not minor technical differences: in clinical decision support, the preferred operating point differs depending on whether the goal is to rule in resistance (prioritizing specificity), avoid withholding an effective drug (prioritizing recall), or balance both. The benchmark therefore makes explicit that no single model dominates all use cases.

### Classical machine learning consistently outperformed deep learning under identical conditions

Neural models, such as MLP, WDNN, CNNGWP, and DeepAMR, were competitive for several tasks, but none consistently surpassed XGBoost or random forest across most drugs and categories. This finding challenges the assumption that increasing architectural complexity necessarily improves translational performance for MTB resistance prediction. It does not imply that deep learning is unsuitable; rather, it suggests that gains depend critically on phenotype quality, feature representation, drug-specific sample size, hyperparameter tuning, and the ability to exploit genomic context beyond sparse variant indicators. Future work should explore whether richer representations (e.g., k-mers, attention over gene sequences, or structure-aware encodings) close this gap.

### Drug difficulty is highly heterogeneous and aligns with clinical urgency

Rifampicin and isoniazid were predicted robustly, consistent with their well-characterized resistance mechanisms and more resistant samples. In contrast, bedaquiline, delamanid, linezolid, and clofazimine remained difficult across models; these drugs are clinically critical for MDR/RR-TB treatment. Sparse resistant examples are a major driver, but not the sole explanation: cycloserine and PAS also had limited support yet achieved higher performance, indicating additional contributions from resistance mechanism complexity, label quality, and feature informativeness. For these emerging drugs, high specificity or accuracy can be misleading if recall and AUPRC remain low. Future benchmarks should prioritise targeted data growth, careful pDST harmonisation, and mechanism-aware features.

### Lineage-aware evaluation reveals incomplete robustness

Under random splits, XGBoost achieved strong performance, but lineage-stratified evaluation showed substantial degradation, particularly in lineage 1 and mixed-lineage settings. This pattern confirms that random splits overestimate real-world robustness when strain population structure, geographic sampling, and resistance mechanisms are unevenly distributed. Before clinical implementation, models should be validated in local populations and periodically re-evaluated as sequencing practices, treatment regimens, and resistance epidemiology change.

### Clinical resistance categories provide an essential translational bridge

RR-TB, MDR-TB, and pan-susceptible were predicted well, whereas **XDR-TB remained a major bottleneck**. This gap reflects both statistical constraints (rare classes provide limited training signal) and biological complexity (composite clinical categories depend on multiple drug-level predictions, each with its own uncertainty). A model that performs well on common first-line drug tasks may therefore still be insufficient for identifying highly resistant or unusual phenotypes. Reporting both drug-level and clinical resistance category metrics is therefore essential for evaluating true translational value. The benchmark’s rule-based category prediction pipeline enables direct assessment of how single-drug errors propagate to higher-level clinical decisions. The benchmark may also facilitate implementation studies in settings where access to rapid phenotypic drug susceptibility testing remains limited.

### Limitations

This benchmark integrates heterogeneous retrospective data; source studies differ in pDST methods, critical concentrations, sequencing protocols, and curation depth. Although we harmonised phenotypes and excluded unreliable records, missing metadata are unlikely to be random, and residual source-level bias cannot be fully eliminated. The benchmark uses retrospective labels and does not include treatment outcomes, prospective clinical decision-making, or cost-effectiveness analysis. Our feature representation emphasizes called single-nucleotide variants and short indels; structural variation, heteroresistance, epistasis, gene expression, and novel resistance mechanisms may require richer encoding. Finally, while we trained all models under identical splits and metrics, deep-learning architectures may benefit from more extensive hyperparameter optimization or larger pre-training; our results should be interpreted as a comparison under fair, not necessarily optimal, conditions for each architecture. Nevertheless, MTB-DRBench provides a reproducible, open, and clinically oriented foundation for future method development, regulatory qualification, and translation of WGS-based resistance prediction into practice.

## 5 Conclusions

MTB-DRBench establishes a large, standardized benchmark for translating WGS-based *M. tuberculosis* drug-resistance prediction into clinically relevant evaluation. Across 18 drug-level tasks and six clinical resistance categories, classical machine-learning models, especially XGBoost, provided the strongest overall performance under identical preprocessing and validation protocols. Persistent weaknesses for emerging drugs and XDR-TB highlight where larger, better-harmonised datasets and mechanism-aware model development are most urgently needed.

## List of abbreviations

AI: artificial intelligence
AMK: amikacin
AMR: antimicrobial resistance
AUPRC: area under the precision–recall curve
AUROC: area under the receiver operating characteristic curve
BDQ: bedaquiline
CAP: capreomycin
CFZ: clofazimine
CNNGWP: convolutional genome-wide prediction
CRyPTIC: Comprehensive Resistance Prediction for Tuberculosis: an International Consortium
CS: cycloserine
DLM: delamanid
DL: deep learning
DR-TB: drug-resistant tuberculosis
DST: drug susceptibility testing
EMB: ethambutol
ETO: ethionamide
F1: F1-score
FQ: fluoroquinolone
HR-TB: isoniazid-resistant tuberculosis
INH: isoniazid
KAN: kanamycin
LFX: levofloxacin
LR: logistic regression
LZD: linezolid
MDR-TB: multidrug-resistant tuberculosis
MFX: moxifloxacin
MIC: minimum inhibitory concentration
ML: machine learning
MLP: multilayer perceptron
MTB: *Mycobacterium tuberculosis*
NPV: negative predictive value
OFX: ofloxacin
PAS: para-aminosalicylic acid
pDST: phenotypic drug susceptibility testing
PPV: positive predictive value
pre-XDR-TB: pre-extensively drug-resistant tuberculosis
PZA: pyrazinamide
RF: random forest
RIF: rifampicin
RR-TB: rifampicin-resistant tuberculosis
SNV: single-nucleotide variant
STM: streptomycin
TB: tuberculosis
WDNN: wide-and-deep neural network
WGS: whole-genome sequencing
XDR-TB: extensively drug-resistant tuberculosis
XGBoost: extreme gradient boosting.

## Additional files

**Additional file 1.** PDF: Supplementary information. Supplementary methods, dataset curation details, additional tables, and supporting benchmark results.

## Declarations

### Ethics approval and consent to participate

Not applicable. This study analysed publicly available, de-identified bacterial whole-genome sequencing and phenotypic drug susceptibility data and involved no participant recruitment, intervention, or identifiable human data; therefore, ethics approval and consent to participate were not required.

### Consent for publication

Not applicable.

### Availability of data and materials

The source whole-genome sequencing and phenotypic drug susceptibility data analysed in this study are available from the public repositories and publications cited in the manuscript. The harmonised benchmark dataset and analysis code are not publicly available at present but are available from the corresponding authors on reasonable request.

### Authors’ contributions

CL and HZ contributed to data curation, development of the bioinformatics pipeline, data preprocessing, and drafting the manuscript. PZ contributed to development of the benchmarking pipeline, interpretation of the results, and drafting the manuscript. NTT, NQD, IA, IHC, ZK, WA, and XZ reviewed and edited the manuscript. HW and YY contributed to study conceptualization, overall supervision, and revision of the manuscript. All authors read and approved the final manuscript.

## Acknowledgements

The computations in this paper were run on the Siyuan-1 cluster supported by the Center for High Performance Computing at Shanghai Jiao Tong University.

