## Supplementary materials for "Benchmarking the translational potential of AI-based drug-resistance prediction from *Mycobacterium tuberculosis* whole-genome sequencing data"

July 3, 2026

#### Supplementary Methods

##### Data curation and source harmonization

The benchmark integrates three sources: CRyPTIC, a published multi-study WGS–pDST compilation, and literature-derived datasets curated from PubMed, Web of Science, and Embase searches. Source identifiers were retained during harmonization to support source-aware auditing and to avoid unintended duplication across public repositories. Isolates without linkable WGS and pDST records were excluded from model-ready analyses.

##### Phenotype harmonization

Drug susceptibility phenotypes were converted to binary susceptible/resistant labels at the isolate–drug level. Missing, indeterminate, contaminated, intermediate, or otherwise uninterpretable phenotypes were excluded from the corresponding drug-specific task. Drug aliases were standardized before model training; for example, capreomycin/CM/CAP, kanamycin/KM/KAN, and streptomycin/SM/STM were mapped to common internal names.

pDST method evidence was additionally grouped into five audit categories: category 1, standard Löwenstein–Jensen, Middlebrook 7H10, Middlebrook 7H11, or BACTEC MGIT testing using WHO critical concentrations; category 2, the same media or platforms with outdated critical concentrations or WHO guidance cited without the tested concentration; category 3, non-standard phenotype methods or interpretive rules not aligned with WHO or CRyPTIC standards; category 4, CRyPTIC-standard Thermo Fisher Scientific broth microdilution MIC testing interpreted using plate-specific epidemiological cutoffs; and category 5, pDST method not reported. These categories were used to describe phenotype-source quality and were not used as model input features.

##### Data splitting and leakage control

Splitting was performed at the isolate level using the `uniqueid` index after duplicate isolates had been resolved. For each drug, eligible isolates were randomly assigned to nested five-fold cross-validation, with folds stratified by the binary drug-resistance label. Each outer iteration allocated 80% of isolates to training and 20% to testing. The outer test fold was not used for model fitting, early stopping, or threshold specification. A further random five-fold split within each outer training set was used for early stopping where applicable. No spatial holdout split was performed. Lineage-stratified results were calculated separately within the lineage groups represented in the outer test folds; lineage labels were not supplied as model features. Phenotypes that were missing or uninterpretable were excluded rather than imputed, and no resampling was applied.

##### Class imbalance and decision thresholds

Class imbalance was addressed within each training fold. Logistic regression (LR) and random forest (RF) used inverse-frequency weighting with the balanced class-weight setting. For XGBoost, the positive-class weight was calculated separately for each drug and fold as

$$\frac{\#\{\text{susceptible isolates in the training fold}\}}{\#\{\text{resistant isolates in the training fold}\}}.$$

The multilayer perceptron (MLP), convolutional genome-wide prediction model (CNNGWP), wide-and-deep neural network (WDNN), and DeepAMR used focal loss with $\gamma = 2.0$. The positive-class weight was adapted within each batch as $\alpha = 1 - \text{positive proportion}$ and clamped to $[0.05, 0.95]$. Predicted probabilities were converted to binary labels using a fixed threshold of 0.5 for every drug and model. This threshold was specified before outer-test evaluation and was not optimized.

#### Metric aggregation and reproducibility

Metrics were calculated independently for each outer test fold. Drug-level values are the mean of five outer folds, and overall model summaries are unweighted macro means across the 18 drug-specific tasks. Error bars report the standard deviation across the five outer folds. When a model made no positive predictions, precision, recall, and F1-score were calculated with `zero_division=0`. AUROC was recorded as undefined, without imputation, when a test fold contained only one class; a drug was omitted from an AUROC summary only when AUROC was undefined in all five folds.

The master random seed was 2026. The same seed was applied to Python, NumPy, scikit-learn (`random_state`), and PyTorch, and deterministic GPU operations were enabled.

#### Model implementations and hyperparameters

**Logistic regression and random forest.** LR and RF were implemented with scikit-learn version 1.8.0. LR used the `LogisticRegression` estimator with an L2 penalty, $C = 0.9$, the `liblinear` solver, balanced class weights, and a maximum of 1000 iterations. RF used `RandomForestClassifier` with 100 trees, unrestricted maximum depth, minimum split and leaf sizes of 2 and 1, respectively, `max_features='sqrt'`, and balanced class weights.

**XGBoost.** XGBoost used `XGBClassifier` from xgboost version 3.1.3, with 1000 boosting rounds, a learning rate of 0.05, maximum depth 6, minimum child weight 1, row and column subsampling fractions of 1.0, `reg_alpha=0`, and `reg_lambda=1`. The class weight was calculated separately in each training fold as described above. Early stopping monitored inner-validation loss with a patience of 30 boosting rounds.

**Neural-network training.** All neural models were implemented in PyTorch version 2.9.1 and optimized with Adam using a learning rate of 0.001, weight decay of $10^{-3}$, and batch size 32. Training was limited to 150 epochs. Early stopping monitored inner-validation loss and was triggered when the loss did not decrease for 15 consecutive epochs. The MLP contained hidden layers of 32 and 16 units, rectified linear unit (ReLU) activations, and dropout of 0.5. Its learning rate was controlled by `ReduceLROnPlateau` in minimization mode with patience 5 and reduction factor 0.5.

The WDNN used an identity wide branch and a deep branch with three 64-unit ReLU layers and dropout of 0.5. The raw input and final 64-dimensional deep representation were concatenated and passed to a linear output layer. CNNGWP treated the binary feature vector as a one-dimensional sequence and applied one convolutional layer with 1 input channel, 16 output channels, kernel size 10, and stride 2, followed by ReLU activation, max pooling with kernel size 2, dropout of 0.5, and a linear output layer.

DeepAMR was configured as a single-drug prediction model. Its encoder widths were 128, 64, and 16 units; the classifier widths were 4 and 1; and the decoder widths were 64, 128, and the input dimension. Dropout of 0.5 provided corruption noise. No separate autoencoder pretraining was performed.

### Supplementary Tables and Files

Table S1: Mapping table for standardizing anti-tuberculosis drug names from multiple sources

| Original Name / Abbreviation | Standardized Full Name | Standardized Abbreviation |
|---|---|---|
| ETH | Ethionamide | ETO |
| OFX | Ofloxacin | OFX |
| PAS | Para-Aminosalicylic Acid | PAS |
| Rifampicin (R) | Rifampicin | RIF |
| tebipenem-clavulanic | Tebipenem-Clavulanic | TEB-CLA |
| Isoniazide | Isoniazid | INH |
| CAP | Capreomycin | CAP |
| Amikacin | Amikacin | AMK |
| RFB | Rifabutin | RFB |
| moxifloxacin | Moxifloxacin | MFX |
| DLM | Delamanid | DLM |
| meropenem-clavulanic | Meropenem-Clavulanic | MER-CLA |
| CIP | Ciprofloxacin | CIP |
| FQs | Fluoroquinolones (group) | FQs |
| PZA | Pyrazinamide | PZA |
| Levofloxacin | Levofloxacin | LFX |
| OFL | Ofloxacin | OFX |
| KM | Kanamycin | KAN |
| Streptomycin | Streptomycin | STM |
| LZD | Linezolid | LZD |
| CFZ | Clofazimine | CFZ |
| ofloxacin | Ofloxacin | OFX |
| Ethambutol | Ethambutol | EMB |
| KAN | Kanamycin | KAN |
| MOX | Moxifloxacin | MFX |
| levofloxacin | Levofloxacin | LFX |
| STM | Streptomycin | STM |
| SM | Streptomycin | STM |
| BDQ | Bedaquiline | BDQ |
| PTH | prothionamide | PTH |
| ETI | Ethionamide | ETO |
| Ethambutol (E) | Ethambutol | EMB |
| pretomanid | Pretomanid | PTM |
| Isoniazid (H) | Isoniazid | INH |
| Rifampicine | Rifampicin | RIF |
| RIF | Rifampicin | RIF |
| Isoniazid | Isoniazid | INH |
| Streptomycin (S) | Streptomycin | STM |
| RFP | Rifampicin | RIF |
| meropenem-vaborbactam | Meropenem-Clavulanic | MER-CLA |
| INH | Isoniazid | INH |
| LVX | Levofloxacin | LFX |
| CS | Cycloserine | CS |
| STR | Streptomycin | STM |
| LFX | Levofloxacin | LFX |
| Kanamycin | Kanamycin | KAN |
| gatifloxacin | Gatifloxacin | GAT |
| RMP | Rifampicin | RIF |
| Pyrazinamide (Z) | Pyrazinamide | PZA |
| AM | Amikacin | AMK |

| Original Name / Abbreviation | Standardized Full Name | Standardized Abbreviation |
|---|---|---|
| EMB | Ethambutol | EMB |
| AK | Amikacin | AMK |
| AMK | Amikacin | AMK |
| ETO | Ethionamide | ETO |
| TH | Ethionamide | ETO |
| CPM | Capreomycin | CAP |
| Moxifloxacin | Moxifloxacin | MFX |
| Rifampicin | Rifampicin | RIF |
| Pyrazinamide | Pyrazinamide | PZA |
| Capreomycin | Capreomycin | CAP |
| MFX | Moxifloxacin | MFX |
| CM | Capreomycin | CAP |

Table S2: Overall model performance summary. Average values of each metric across all 18 drugs for seven methods (LR, RF, XGB, MLP, CNNGWP, WDNN, DeepAMR). Best values per metric are shown in **bold**.

| **Metric** | **LR** | **RF** | **XGB** | **MLP** | **CNNGWP** | **WDNN** | **DeepAMR** |
|---|---|---|---|---|---|---|---|
| AUROC | 0.8803 | 0.8726 | 0.8712 | 0.8868 | 0.8788 | **0.8906** | 0.8095 |
| AUPRC | 0.6443 | 0.6457 | **0.6741** | 0.6572 | 0.6273 | 0.6551 | 0.5850 |
| Sensitivity | **0.7737** | 0.6291 | 0.7222 | 0.7051 | 0.6745 | 0.6977 | 0.6457 |
| Specificity | 0.8919 | **0.9565** | 0.9351 | 0.9342 | 0.9258 | 0.9364 | 0.8891 |
| Accuracy | 0.8844 | **0.9325** | 0.9229 | 0.9258 | 0.9158 | 0.9263 | 0.8849 |
| Precision | 0.5140 | **0.6151** | 0.5757 | 0.5443 | 0.5204 | 0.5477 | 0.4274 |
| F1 | 0.5809 | 0.6107 | **0.6203** | 0.6057 | 0.5712 | 0.6037 | 0.5004 |
| PPV | 0.5140 | **0.6151** | 0.5757 | 0.5443 | 0.5204 | 0.5477 | 0.4274 |
| NPV | 0.9747 | 0.9637 | 0.9744 | **0.9757** | 0.9714 | 0.9743 | 0.9660 |

Table S3: AUROC scores of seven methods across 18 drugs (overall). Values are shown as mean ± standard deviation across five outer folds. Best values per drug are shown in **bold**.

| **Drug** | **LR** | **RF** | **XGB** | **MLP** | **CNNGWP** | **WDNN** | **DeepAMR** | **Best** |
|---|---|---|---|---|---|---|---|---|
| AMK | 0.9296±0.006 | 0.9233±0.005 | 0.9208±0.006 | **0.9301±0.007** | 0.9236±0.008 | 0.9300±0.008 | 0.9058±0.003 | MLP |
| BDQ | 0.7027±0.056 | 0.6520±0.048 | 0.5918±0.037 | 0.7172±0.074 | 0.7342±0.076 | **0.7561±0.078** | 0.4560±0.083 | WDNN |
| CAP | 0.9122±0.006 | 0.9148±0.012 | 0.9107±0.002 | **0.9202±0.003** | 0.9078±0.004 | 0.9170±0.005 | 0.9001±0.007 | MLP |
| CFZ | 0.7654±0.025 | 0.6909±0.040 | 0.7126±0.028 | 0.7804±0.016 | 0.7832±0.016 | **0.7900±0.014** | 0.5691±0.025 | WDNN |
| CS | 0.8958±0.010 | **0.9521±0.012** | 0.9408±0.016 | 0.9195±0.011 | 0.8848±0.015 | 0.9236±0.009 | 0.7353±0.135 | RF |
| DLM | **0.6690±0.041** | 0.6309±0.036 | 0.6099±0.074 | 0.6392±0.029 | 0.6252±0.040 | 0.6670±0.030 | 0.4778±0.062 | LR |
| EMB | 0.9632±0.003 | 0.9592±0.003 | 0.9649±0.003 | 0.9640±0.002 | 0.9564±0.003 | **0.9652±0.002** | 0.9578±0.003 | WDNN |
| ETO | 0.8852±0.010 | 0.8722±0.013 | 0.8907±0.011 | **0.8908±0.012** | 0.8751±0.013 | 0.8894±0.010 | 0.8707±0.012 | MLP |
| INH | 0.9552±0.001 | 0.9545±0.001 | **0.9570±0.002** | 0.9559±0.001 | 0.9502±0.002 | 0.9561±0.002 | 0.9517±0.003 | XGB |
| KAN | 0.9106±0.008 | 0.8874±0.011 | 0.9060±0.008 | **0.9151±0.006** | 0.9033±0.009 | 0.9126±0.008 | 0.8972±0.008 | MLP |
| LFX | 0.9462±0.006 | 0.9379±0.007 | 0.9462±0.004 | 0.9475±0.005 | 0.9447±0.004 | **0.9484±0.004** | 0.9356±0.005 | WDNN |
| LZD | 0.6481±0.064 | 0.6858±0.067 | 0.6355±0.073 | 0.7032±0.058 | **0.7116±0.061** | 0.7101±0.055 | 0.4234±0.061 | CNNGWP |
| MFX | 0.9351±0.010 | 0.9179±0.012 | 0.9303±0.012 | 0.9382±0.008 | 0.9323±0.008 | **0.9396±0.008** | 0.9162±0.008 | WDNN |
| OFX | 0.9504±0.012 | 0.9548±0.016 | **0.9584±0.009** | 0.9543±0.010 | 0.9496±0.010 | 0.9555±0.010 | 0.9385±0.015 | XGB |
| PAS | 0.8786±0.045 | 0.8832±0.031 | **0.9002±0.036** | 0.8850±0.049 | 0.8527±0.051 | 0.8688±0.045 | 0.7596±0.106 | XGB |
| PZA | 0.9594±0.005 | 0.9566±0.007 | **0.9631±0.006** | 0.9616±0.006 | 0.9547±0.004 | 0.9610±0.005 | 0.9525±0.005 | XGB |
| RIF | 0.9796±0.002 | 0.9782±0.002 | 0.9806±0.002 | 0.9806±0.002 | 0.9758±0.003 | **0.9807±0.002** | 0.9757±0.002 | WDNN |
| STM | 0.9589±0.003 | 0.9545±0.002 | **0.9617±0.002** | 0.9595±0.003 | 0.9526±0.002 | 0.9598±0.004 | 0.9483±0.002 | XGB |

Table S4: Best performing method for each drug across all metrics.

| Drug | AUROC | AUPRC | Sensitivity | Specificity | Accuracy | Precision | F1-score | PPV | NPV |
|---|---|---|---|---|---|---|---|---|---|
| AMK | MLP(0.9301) | XGB(0.8140) | DeepAMR(0.8338) | RF(0.9857) | RF(0.9695) | RF(0.7917) | RF(0.7676) | RF(0.7917) | LR(0.9862) |
| BDQ | WDNN(0.7561) | WDNN(0.1028) | LR(0.4333) | CNNGWP(1.0000) | CNNGWP(0.9922) | WDNN(0.1030) | WDNN(0.0860) | WDNN(0.1030) | LR(0.9949) |
| CAP | MLP(0.9202) | XGB(0.7698) | DeepAMR(0.8206) | RF(0.9723) | RF(0.9403) | RF(0.7474) | XGB(0.7132) | RF(0.7474) | MLP(0.9747) |
| CFZ | WDNN(0.7900) | WDNN(0.1079) | LR(0.6364) | DeepAMR(1.0000) | DeepAMR(0.9800) | CNNGWP(0.1159) | WDNN(0.1642) | CNNGWP(0.1159) | LR(0.9905) |
| CS | RF(0.9521) | RF(0.9095) | XGB(0.9053) | RF(0.9132) | XGB(0.8961) | RF(0.8075) | XGB(0.8375) | RF(0.8075) | XGB(0.9575) |
| DLM | LR(0.6690) | XGB(0.0845) | LR(0.4993) | CNNGWP(1.0000) | CNNGWP(0.9873) | WDNN(0.0867) | XGB(0.0585) | WDNN(0.0867) | LR(0.9917) |
| EMB | WDNN(0.9652) | XGB(0.8579) | DeepAMR(0.9312) | RF(0.9663) | RF(0.9389) | RF(0.7852) | XGB(0.7930) | RF(0.7852) | DeepAMR(0.9880) |
| ETO | MLP(0.8908) | XGB(0.7049) | LR(0.7835) | RF(0.9492) | RF(0.8992) | RF(0.6505) | XGB(0.6443) | RF(0.6505) | LR(0.9608) |
| INH | XGB(0.9570) | XGB(0.9427) | WDNN(0.8821) | CNNGWP(0.9787) | XGB(0.9447) | CNNGWP(0.9448) | XGB(0.9046) | CNNGWP(0.9448) | WDNN(0.9505) |
| KAN | MLP(0.9151) | MLP(0.7672) | LR(0.7956) | RF(0.9486) | RF(0.9203) | RF(0.5952) | RF(0.6298) | RF(0.5952) | LR(0.9746) |
| LFX | WDNN(0.9484) | WDNN(0.8658) | MLP(0.8665) | RF(0.9734) | XGB(0.9468) | RF(0.8312) | XGB(0.8240) | RF(0.8312) | MLP(0.9769) |
| LZD | CNNGWP(0.7116) | MLP(0.1922) | LR(0.4301) | DeepAMR(1.0000) | DeepAMR(0.9892) | CNNGWP(0.4036) | MLP(0.2583) | CNNGWP(0.4036) | LR(0.9926) |
| MFX | WDNN(0.9396) | MLP(0.7323) | MLP(0.8841) | RF(0.9666) | RF(0.9311) | RF(0.6719) | XGB(0.6953) | RF(0.6719) | MLP(0.9861) |
| OFX | XGB(0.9584) | XGB(0.8973) | CNNGWP(0.8720) | RF(0.9698) | XGB(0.9437) | RF(0.8326) | XGB(0.8333) | RF(0.8326) | CNNGWP(0.9725) |
| PAS | XGB(0.9002) | XGB(0.7007) | MLP(0.8227) | RF(0.9468) | RF(0.9201) | RF(0.6136) | RF(0.6415) | RF(0.6136) | MLP(0.9766) |
| PZA | XGB(0.9631) | XGB(0.8906) | MLP(0.8966) | RF(0.9679) | XGB(0.9457) | RF(0.8315) | XGB(0.8404) | RF(0.8315) | MLP(0.9791) |
| RIF | WDNN(0.9807) | XGB(0.9625) | WDNN(0.9419) | RF(0.9829) | XGB(0.9722) | XGB(0.9437) | XGB(0.9410) | XGB(0.9437) | WDNN(0.9819) |
| STM | XGB(0.9617) | XGB(0.9368) | MLP(0.9082) | RF(0.9344) | XGB(0.9141) | RF(0.8690) | XGB(0.8757) | RF(0.8690) | MLP(0.9515) |

Table S5: Clinical resistance category performance summary. Average metric values across seven methods for each clinical resistance category. Sample sizes: RR-TB (n=47,780), MDR-TB (n=46,416), pre-XDR-TB (n=25,444), XDR-TB (n=12,686), HR-TB (n=46,416), Pan-Susceptible (n=14,755).

| Metric | RR-TB | MDR-TB | pre-XDR | XDR-TB | HR-TB | Pan-Susc. |
|---|---|---|---|---|---|---|
| Sensitivity | 0.9323 | 0.9349 | 0.8755 | 0.3019 | 0.5257 | 0.9656 |
| Specificity | 0.9786 | 0.9760 | 0.9531 | 0.9880 | 0.9866 | 0.8635 |
| Accuracy | 0.9677 | 0.9671 | 0.9443 | 0.9828 | 0.9508 | 0.9314 |
| Precision | 0.9316 | 0.9164 | 0.7299 | 0.2241 | 0.7695 | 0.9335 |
| F1-score | 0.9319 | 0.9253 | 0.7888 | 0.2144 | 0.6200 | 0.9492 |
| PPV | 0.9316 | 0.9164 | 0.7299 | 0.2241 | 0.7695 | 0.9335 |
| NPV | 0.9790 | 0.9818 | 0.9836 | 0.9946 | 0.9612 | 0.9272 |

Table S6: AUROC scores of seven methods across 18 drugs and five *M. tuberculosis* lineages. Values are shown as mean ± standard deviation across five outer folds. Best values per lineage–drug combination are in **bold**. '—' indicates insufficient samples to compute the metric.

| Drug | Lineage | LR | RF | XGB | MLP | CNNGWP | WDNN | DeepAMR |
|---|---|---|---|---|---|---|---|---|
| AMK | L1 | **0.8660±0.055** | 0.8496±0.113 | 0.7751±0.097 | 0.7911±0.098 | 0.8546±0.069 | 0.7024±0.088 | 0.7043±0.180 |
| | L2 | 0.9291±0.006 | 0.9217±0.011 | 0.9180±0.012 | 0.9375±0.013 | 0.9379±0.012 | **0.9389±0.010** | 0.8473±0.195 |
| | L3 | 0.9457±0.016 | **0.9705±0.018** | 0.9358±0.028 | 0.9467±0.040 | 0.9585±0.031 | 0.9529±0.013 | 0.7779±0.254 |
| | L4 | 0.9061±0.011 | 0.9039±0.021 | 0.8919±0.018 | 0.9085±0.016 | 0.9047±0.022 | **0.9115±0.017** | 0.8115±0.175 |
| | Mixed | 0.6908±0.052 | 0.7274±0.102 | 0.7427±0.128 | 0.6590±0.119 | **0.7551±0.167** | 0.6859±0.187 | 0.6526±0.040 |
| BDQ | L2 | 0.6949±0.108 | 0.6544±0.079 | 0.5474±0.134 | 0.7139±0.093 | **0.7978±0.069** | 0.7200±0.102 | 0.6624±0.122 |
| | L3 | 0.7941±0.073 | 0.8048±0.100 | **0.8093±0.065** | 0.5901±0.177 | 0.7222±0.102 | 0.6908±0.154 | 0.5767±0.190 |
| | L4 | 0.5234±0.144 | 0.5178±0.073 | 0.5228±0.205 | 0.5758±0.083 | 0.5831±0.207 | 0.5736±0.135 | **0.5930±0.169** |
| CAP | L1 | 0.7659±0.094 | 0.7508±0.087 | **0.8205±0.100** | 0.6466±0.197 | 0.6158±0.124 | 0.6319±0.219 | 0.5366±0.112 |
| | L2 | 0.9067±0.013 | 0.9030±0.019 | 0.9062±0.016 | **0.9109±0.012** | 0.9082±0.018 | 0.9073±0.018 | 0.9007±0.018 |
| | L3 | 0.7695±0.071 | **0.8656±0.072** | 0.8428±0.034 | 0.8212±0.076 | 0.8489±0.074 | 0.7789±0.044 | 0.7812±0.151 |
| | L4 | 0.9210±0.019 | **0.9217±0.018** | 0.9100±0.019 | 0.9213±0.017 | 0.9137±0.015 | 0.9208±0.019 | 0.9175±0.013 |
| | Mixed | 0.8423±0.133 | **0.9013±0.133** | 0.8603±0.140 | 0.8462±0.133 | 0.8269±0.119 | 0.8744±0.142 | 0.7987±0.130 |
| CFZ | L1 | 0.3605±0.109 | 0.4800±0.005 | **0.6729±0.243** | — | — | — | — |
| | L2 | 0.7361±0.030 | 0.6502±0.033 | 0.6631±0.042 | 0.7573±0.017 | **0.7770±0.027** | 0.7607±0.025 | 0.6918±0.045 |
| | L3 | **0.6236±0.251** | 0.5565±0.233 | 0.6182±0.168 | 0.4914±0.252 | 0.2434±0.160 | 0.3710±0.356 | 0.4357±0.264 |
| | L4 | 0.6928±0.090 | 0.6386±0.093 | 0.6336±0.144 | 0.7222±0.103 | **0.7265±0.045** | 0.7069±0.077 | 0.6466±0.135 |
| | Mixed | 0.7658±0.162 | **0.7898±0.208** | 0.7640±0.219 | 0.7500±0.317 | 0.5263±0.077 | 0.6307±0.282 | 0.6953±0.204 |
| CS | L2 | 0.8999±0.050 | **0.9571±0.028** | 0.9563±0.044 | 0.9354±0.039 | 0.8808±0.073 | 0.9257±0.053 | 0.8755±0.061 |
| | L4 | 0.9128±0.041 | 0.9233±0.043 | **0.9392±0.016** | 0.9121±0.043 | 0.8928±0.056 | 0.9015±0.049 | 0.8856±0.058 |
| DLM | L1 | 0.8449±0.102 | 0.6932±0.282 | **0.8828±0.105** | 0.8619±0.054 | 0.7512±0.347 | 0.6450±0.233 | 0.7739±0.114 |
| | L2 | **0.6734±0.080** | 0.6366±0.072 | 0.6144±0.074 | 0.6317±0.047 | 0.6197±0.061 | 0.6485±0.058 | 0.6017±0.029 |
| | L3 | 0.4686±0.276 | **0.6439±0.239** | 0.6260±0.259 | 0.4691±0.266 | 0.2559±0.152 | 0.3375±0.300 | 0.4253±0.301 |
| | L4 | **0.6650±0.051** | 0.5746±0.103 | 0.5772±0.066 | 0.6344±0.120 | 0.6037±0.066 | 0.6215±0.050 | 0.6151±0.103 |
| | Mixed | **0.6140±0.110** | 0.5175±0.312 | 0.2982±0.171 | — | — | — | — |
| EMB | L1 | 0.9370±0.017 | 0.9395±0.011 | 0.9300±0.025 | 0.9315±0.015 | **0.9431±0.024** | 0.9398±0.023 | 0.7526±0.232 |
| | L2 | 0.9584±0.001 | 0.9553±0.002 | **0.9626±0.001** | 0.9608±0.002 | 0.9543±0.001 | 0.9587±0.001 | 0.9522±0.001 |
| | L3 | 0.9582±0.013 | 0.9595±0.006 | 0.9602±0.010 | **0.9606±0.008** | 0.9548±0.009 | 0.9571±0.008 | 0.8685±0.206 |
| | L4 | 0.9597±0.006 | 0.9546±0.010 | 0.9588±0.006 | **0.9612±0.009** | 0.9538±0.006 | 0.9597±0.008 | 0.9555±0.006 |
| | Mixed | 0.8343±0.152 | **0.9093±0.064** | 0.8343±0.107 | 0.7890±0.137 | 0.7799±0.144 | 0.7906±0.130 | 0.7998±0.209 |
| ETO | L1 | 0.8273±0.044 | 0.8247±0.035 | 0.8292±0.039 | **0.8447±0.042** | 0.8389±0.035 | 0.8386±0.037 | 0.8318±0.021 |
| | L2 | 0.8999±0.008 | 0.8856±0.010 | **0.9041±0.010** | 0.9011±0.009 | 0.8948±0.010 | 0.9014±0.008 | 0.8835±0.008 |
| | L3 | 0.7988±0.058 | 0.7459±0.043 | 0.7455±0.061 | 0.8015±0.059 | 0.8082±0.069 | **0.8133±0.047** | 0.7491±0.129 |
| | L4 | 0.8723±0.017 | 0.8644±0.010 | 0.8690±0.010 | 0.8747±0.023 | 0.8629±0.013 | **0.8757±0.014** | 0.7586±0.238 |
| | Mixed | 0.6890±0.150 | **0.7694±0.130** | 0.6790±0.163 | 0.7099±0.090 | 0.7264±0.128 | 0.5953±0.114 | 0.6278±0.133 |
| INH | L1 | **0.9268±0.015** | 0.9038±0.019 | 0.9148±0.019 | 0.9191±0.024 | 0.9230±0.015 | 0.9244±0.019 | 0.8315±0.186 |
| | L2 | 0.9702±0.003 | 0.9690±0.002 | 0.9705±0.002 | 0.9697±0.003 | 0.9691±0.003 | **0.9710±0.003** | 0.9698±0.003 |
| | L3 | 0.9499±0.008 | 0.9487±0.010 | 0.9520±0.009 | 0.9533±0.007 | **0.9558±0.007** | 0.9512±0.007 | 0.9539±0.006 |
| | L4 | 0.9412±0.003 | 0.9424±0.003 | **0.9442±0.003** | 0.9422±0.004 | 0.9369±0.003 | 0.9421±0.004 | 0.9373±0.005 |

| Drug | Lineage | LR | RF | XGB | MLP | CNNGWP | WDNN | DeepAMR |
|---|---|---|---|---|---|---|---|---|
| | Mixed | 0.8773±0.043 | **0.8799±0.032** | 0.8781±0.036 | 0.8551±0.053 | 0.8616±0.029 | 0.8564±0.058 | 0.8488±0.056 |
| KAN | L1 | 0.8370±0.108 | 0.8070±0.072 | 0.7829±0.096 | **0.8392±0.081** | 0.7966±0.062 | 0.8290±0.080 | 0.6305±0.227 |
| | L2 | 0.9324±0.014 | 0.9197±0.006 | 0.9215±0.010 | **0.9354±0.014** | 0.9341±0.013 | 0.9337±0.012 | 0.9259±0.016 |
| | L3 | 0.9007±0.042 | 0.9100±0.044 | 0.8733±0.038 | 0.9037±0.050 | **0.9218±0.041** | 0.9153±0.051 | 0.7914±0.177 |
| | L4 | 0.8708±0.015 | 0.8499±0.010 | 0.8623±0.013 | 0.8730±0.016 | 0.8641±0.022 | **0.8748±0.018** | 0.7489±0.160 |
| | Mixed | 0.6214±0.061 | 0.6218±0.059 | 0.5864±0.113 | 0.5959±0.077 | 0.6295±0.100 | **0.6559±0.061** | 0.5886±0.188 |
| LFX | L1 | 0.8222±0.063 | 0.8195±0.081 | 0.8172±0.099 | 0.7958±0.057 | **0.8439±0.057** | 0.6988±0.111 | 0.7471±0.175 |
| | L2 | 0.9592±0.007 | 0.9533±0.006 | 0.9609±0.006 | 0.9600±0.006 | 0.9598±0.007 | **0.9617±0.008** | 0.9488±0.006 |
| | L3 | **0.9368±0.020** | 0.9225±0.016 | 0.9324±0.015 | 0.9331±0.016 | 0.9333±0.022 | 0.9345±0.016 | 0.8420±0.192 |
| | L4 | 0.9153±0.017 | 0.8982±0.003 | 0.8972±0.019 | 0.9148±0.015 | 0.9143±0.013 | **0.9193±0.014** | 0.6607±0.220 |
| | Mixed | **0.6828±0.066** | 0.6789±0.042 | 0.6620±0.082 | 0.6196±0.094 | 0.6055±0.067 | 0.6329±0.091 | 0.5975±0.117 |
| LZD | L1 | 0.2283±0.195 | 0.4110±0.023 | 0.4529±0.246 | 0.2776±0.313 | 0.3140±0.391 | 0.3796±0.385 | **0.5229±0.284** |
| | L2 | 0.6692±0.087 | 0.6828±0.046 | 0.6214±0.093 | 0.7164±0.123 | 0.7446±0.094 | **0.7486±0.101** | 0.5042±0.118 |
| | L3 | 0.4212±0.197 | 0.6603±0.231 | **0.7219±0.158** | 0.5732±0.121 | 0.3635±0.270 | 0.5513±0.314 | 0.3961±0.179 |
| | L4 | 0.6059±0.150 | 0.6316±0.092 | 0.6346±0.158 | 0.6241±0.153 | **0.6833±0.132** | 0.5990±0.139 | 0.6245±0.127 |
| | Mixed | 0.2828±0.282 | **0.4949±0.313** | 0.4697±0.427 | — | — | — | — |
| MFX | L1 | 0.7273±0.153 | 0.7830±0.119 | 0.7415±0.115 | 0.7439±0.135 | **0.8092±0.112** | 0.7644±0.144 | 0.7292±0.149 |
| | L2 | 0.9416±0.004 | 0.9282±0.008 | **0.9440±0.003** | 0.9438±0.002 | 0.9374±0.003 | 0.9431±0.005 | 0.9221±0.004 |
| | L3 | 0.8850±0.058 | 0.8890±0.023 | 0.8686±0.059 | 0.8745±0.045 | **0.8910±0.047** | 0.8761±0.040 | 0.7309±0.146 |
| | L4 | 0.8949±0.018 | 0.8713±0.010 | 0.8753±0.008 | 0.8978±0.018 | 0.9013±0.023 | **0.9032±0.024** | 0.7405±0.221 |
| | Mixed | 0.6918±0.203 | 0.6794±0.073 | 0.6137±0.084 | 0.5234±0.122 | **0.7059±0.153** | 0.6205±0.238 | 0.6545±0.170 |
| OFX | L1 | 0.7991±0.199 | **0.8582±0.090** | 0.8454±0.182 | 0.7286±0.175 | 0.8030±0.110 | 0.7692±0.135 | 0.6819±0.080 |
| | L2 | 0.9778±0.003 | 0.9758±0.005 | **0.9784±0.002** | 0.9770±0.006 | 0.9752±0.003 | 0.9755±0.006 | 0.8684±0.206 |
| | L3 | 0.9355±0.022 | **0.9477±0.021** | 0.9426±0.014 | 0.9080±0.051 | 0.9358±0.027 | 0.9225±0.028 | 0.7955±0.206 |
| | L4 | 0.9260±0.041 | 0.9280±0.027 | 0.9250±0.031 | **0.9314±0.025** | 0.9251±0.024 | 0.9245±0.031 | 0.8429±0.192 |
| | Mixed | **0.8208±0.243** | 0.7333±0.435 | 0.7903±0.309 | 0.7694±0.225 | 0.7542±0.313 | 0.7556±0.238 | 0.7333±0.253 |
| PAS | L2 | 0.8087±0.101 | 0.8626±0.032 | **0.8628±0.055** | 0.7925±0.091 | 0.8039±0.073 | 0.7857±0.086 | 0.6596±0.105 |
| | L4 | 0.9295±0.024 | 0.9301±0.034 | **0.9325±0.024** | 0.9245±0.028 | 0.8888±0.040 | 0.8881±0.051 | 0.8563±0.070 |
| PZA | L1 | 0.7859±0.047 | 0.7784±0.026 | **0.7940±0.038** | 0.7824±0.025 | 0.7928±0.039 | 0.7799±0.034 | 0.7860±0.044 |
| | L2 | 0.9440±0.005 | 0.9463±0.005 | **0.9523±0.002** | 0.9456±0.005 | 0.9407±0.007 | 0.9433±0.007 | 0.9314±0.010 |
| | L3 | 0.9742±0.015 | 0.9847±0.013 | **0.9866±0.006** | 0.9843±0.008 | 0.9832±0.007 | 0.9808±0.012 | 0.8900±0.218 |
| | L4 | 0.9577±0.008 | **0.9605±0.009** | 0.9602±0.007 | 0.9565±0.008 | 0.9530±0.009 | 0.9587±0.010 | 0.8604±0.202 |
| | Mixed | 0.9708±0.033 | 0.9773±0.025 | **0.9788±0.019** | 0.9657±0.036 | 0.9660±0.030 | 0.9668±0.034 | 0.9527±0.039 |
| RIF | L1 | **0.9667±0.013** | 0.9493±0.016 | 0.9610±0.011 | 0.9641±0.016 | 0.9644±0.014 | 0.9658±0.009 | 0.9643±0.010 |
| | L2 | 0.9815±0.003 | 0.9811±0.003 | 0.9827±0.002 | **0.9832±0.002** | 0.9812±0.003 | 0.9822±0.002 | 0.9794±0.002 |
| | L3 | 0.9851±0.004 | 0.9807±0.003 | **0.9864±0.005** | 0.9842±0.006 | 0.9831±0.003 | 0.9843±0.005 | 0.9812±0.004 |
| | L4 | 0.9734±0.006 | 0.9723±0.008 | **0.9745±0.006** | 0.9739±0.007 | 0.9701±0.007 | 0.9738±0.006 | 0.9695±0.005 |
| | Mixed | 0.8899±0.056 | **0.9073±0.033** | 0.9009±0.042 | 0.8784±0.046 | 0.9032±0.035 | 0.8940±0.037 | 0.8692±0.059 |
| STM | L1 | 0.9195±0.015 | 0.9238±0.021 | **0.9258±0.030** | 0.9211±0.028 | 0.9133±0.034 | 0.9194±0.017 | 0.8380±0.079 |
| | L2 | **0.9754±0.005** | 0.9711±0.006 | 0.9737±0.004 | 0.9737±0.003 | 0.9732±0.005 | 0.9733±0.004 | 0.9689±0.005 |
| | L3 | 0.9216±0.014 | 0.9184±0.018 | **0.9264±0.013** | 0.9254±0.018 | 0.9249±0.018 | 0.9208±0.018 | 0.9155±0.022 |
| | L4 | 0.9318±0.004 | 0.9305±0.005 | **0.9371±0.004** | 0.9345±0.004 | 0.9283±0.005 | 0.9342±0.003 | 0.8410±0.191 |
| | Mixed | 0.9419±0.052 | 0.9325±0.067 | 0.9332±0.055 | 0.9339±0.064 | **0.9554±0.025** | 0.9457±0.048 | 0.8478±0.198 |
